# Developing single molecule methods for measuring the pathway proteins ERK, AKT, cyclin d and p70s6k in localized colon cancer in relation to mutation status

**DOI:** 10.1101/695809

**Authors:** Dorte Aa. Olsen, Caroline EB. Thomsen, Rikke F. Andersen, Jonna S. Madsen, Anders Jakobsen, Ivan Brandslund

## Abstract

**Background:** The aim of this study was to quantify the intracellular pathway proteins ERK, AKT, cyclin d and p70s6k in localized colon cancer tissue to investigate the possible prognostic values and the ability to be used as screening markers for upstream mutations.

**Methods:** Colon cancer tissue and autologous reference tissue were collected from 176 patients who underwent surgery for colon cancer. Assays for quantifying ERK, AKT, cyclin d and p70s6k proteins were developed using single molecule array (Simoa). KRAS/BRAF/PIK3CA mutation status was determined using droplet digital PCR.

**Results:** Patients with BRAF mutations had decreased concentrations of ERK (p=0.0002), AKT (p=0.00004) and cyclin d (p=0.001) while no significant differences were found between patients with KRAS mutations and Wild type (Wt) patients. None of the investigated protein concentrations were associated with disease free survival or overall survival, if including all patients. However, when stratifying according to mutation status, significant correlations to overall survival were seen for patients with BRAF mutations and AKT (p=0.003) or ERK (p=0.046) and for patients with KRAS mutations and p70s6k (p=0.04). Furthermore, the combination of genetic mutations, stage 2 disease, and all of the investigated pathway proteins showed significant correlations to overall survival.

**Conclusions:** There is a strong correlation between pathway protein concentrations and mutational BRAF status. Overall survival in colon cancer patients depend both on gene mutation status and pathway protein concentrations. As significant correlations were found between BRAF mutations and ERK, AKT and cyclin d, concentration measurements of these pathway proteins might be useful as screening for upstream mutations.

## Introduction

The intracellular signaling network of the epidermal growth factor receptor (EGFr) consists of two key signal pathways. The mitogen-activated protein kinase (MAPK) also termed RAS/RAF/MEK/ERK and the phosphatidylinositol 3-kinase (PI3K)/ protein kinase B (AKT) pathways. They interact in a complex coordinated manner to regulate all stimulated cellular processes and have been described in detail [1;2]. Both ERK and AKT activate more than 100 downstream proteins from the cytosol to the nucleus, including transcription factors, protein kinases, phosphatases and cytoskeletal elements. Thus, they are involved in a wide variety of nuclear and cytosolic processes including cell differentiation, proliferation and oncogenic transformation [3–5]. One important substrate downstream of AKT is mammalian target of rapamycin (mTOR), which promotes protein translation and cell growth through the activation of p70 ribosomal protein S6 kinase (p70s6k) and the eukaryotic translation initiation factor 4E-binding protein 1 (4EBP1) [6]. Upon activation synthesis of cyclin d increases, leading to protein synthesis, cell growth and cell cycle progression. A schematic overview of the pathways is illustrated in figure 1.

**Fig 1.**
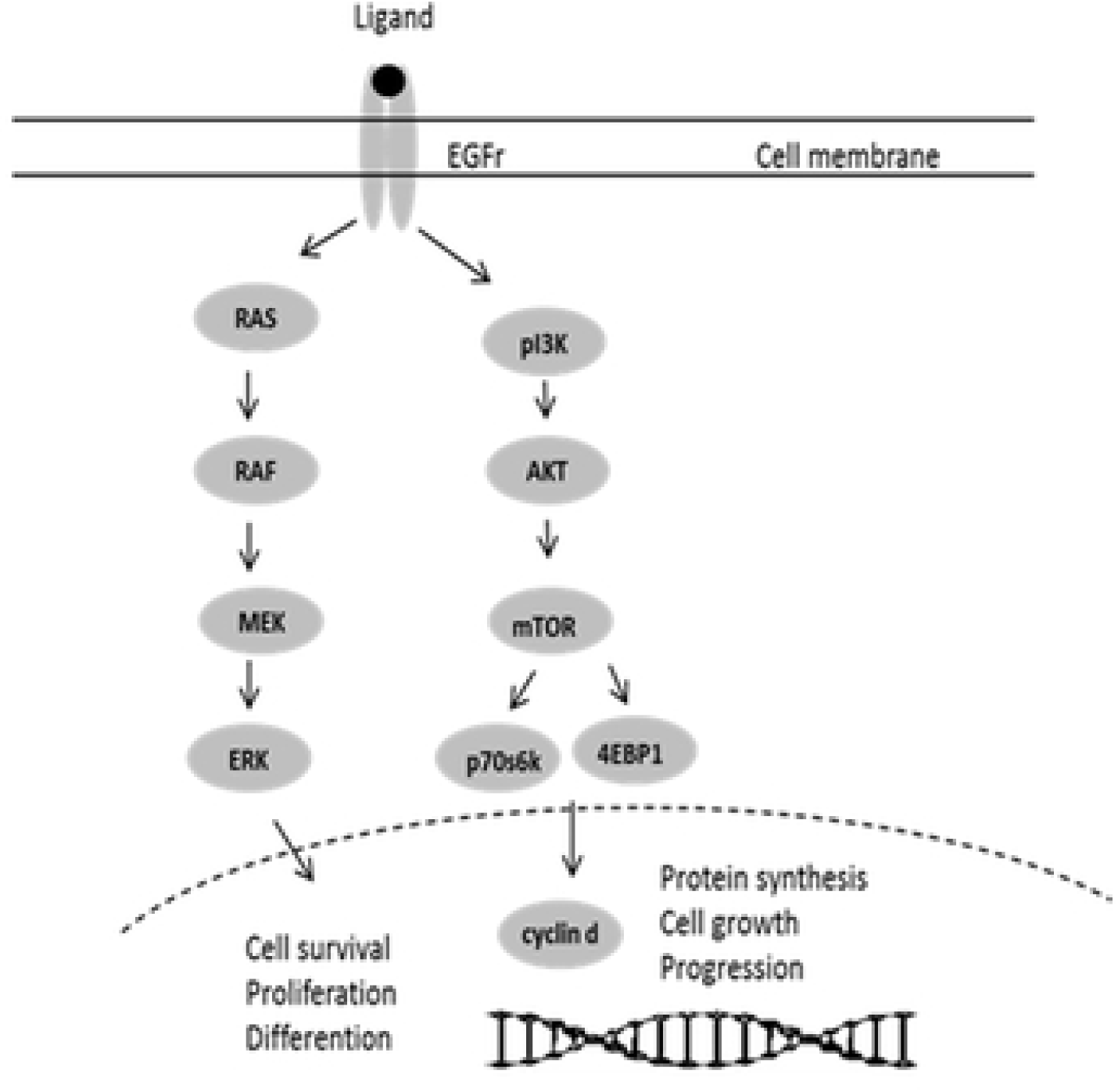
Simplified graphic illustration of the EGFr pathways RAS/RAF/MEK/ERK and PI3K/AKT.

Activation and dysregulation of intracellular signaling pathways plays a critical role in cancer. A frequent alteration in signaling in colorectal cancer is in the RAS and RAF proteins which result in the proteins being constitutively active and stimulating the ERK signaling pathway even though no signal is present. The occurrence of KRAS and BRAF mutations in colorectal cancers has been found to be approximately 40% and 10-25%, respectively [7–9]. Also dysregulation in the PI3K/AKT pathway due to activating mutations in PI3K (PIK3CA) has been identified in colorectal cancer [10–12] and PIK3CA mutations have been found to commonly coexist with KRAS or BRAF mutations [11].

The use of inhibitors against growth factor receptors and tyrosine kinase activators has become standard anti- cancer therapy during the last 10 to 15 years. Some of these monoclonal antibodies used in the treatment are Cetuximab as a blocker to EGFr in colorectal cancer and the Trastuzumab HER2 receptor blocker in breast cancer. Mutations in the receptor proteins or in the pathway proteins result in resistance to treatment using these monoclonal antibodies. It is therefore important to detect such mutations at an early phase before treating the patients as it can be predicted whether the treatment will be without effect and thus the patients will only experience side effects of the treatment.

Usually known mutations are diagnosed using PCR or sequencing which are laborious, time consuming, expensive and causes a delay of several days for reporting. In the future, the number of mutations in the pathway proteins will increase and hence will the expense for sequencing or detecting mutations by PCR. Alternative methodology to detect activation of the intracellular pathway proteins could gain importance especially if such methods would be both faster and considerable cheaper.

We therefore aimed to see whether quantification of pathway proteins in colon cancer tissue might reflect upstream mutations and to detect whether changes in concentrations might be correlated to effect of treatment or clinical outcome. We developed quantitative protein assays for measuring phosphorylated ERK (pERK) as a marker of MAPK pathway activation, phosphorylated AKT (pAKT) for PI13K/AKT pathway activation and phosphorylated p70s6k (pp70s6k) for mTOR activation. Also the total protein levels of ERK (tERK), AKT (tAKT), and cyclin d were measured.

## Materials and methods

### Patients

A total of 176 patients who underwent surgery for colon cancer at Vejle Hospital during 2010-13 were included in the study. Patients with TNM stage 1 (n=8), stage 2 (n=96), and stage 3 (n=68). Four patients had no clinical data available. The study was approved by the Ethics Committee for Southern Denmark (S-20140178).

### Samples and controls

Colon cancer tissue was dissected together with autologous reference tissue by an experienced pathologist. The tissue was stored in RNAlater (Qiagen, Hilden, Germany) at −20°C until use. Colon cancer tissue and autologous reference tissue were homogenized in lysis buffer (50mM Tris-HCL, 150 mM NaCl, pH 7.5, 1% triton X-100) including protease and phosphatase inhibitor cocktail 10µl/ml lysis buffer using the dispomix system (Xiril, Hombrechtikon, Switzerland). The samples were afterwards centrifuged at 16.000 g at 4°C for 15 minutes and the supernatant were recovered, aliquoted and stored at −80°C until use. The total protein concentration was measured using the BCA protein assay reagent (Pierce, Rockford, IL, USA). The tissue lysate samples were diluted in lysis buffer to a final concentration of 1 mg/ml and afterwards diluted further in the specific assay reagent. Matched colon cancer tissue and autologous reference tissue samples were analyzed in the same run. Two controls were prepared using reference colon tissue treated equal to the samples. The controls were included in each run and used to determine intra-assay and total coefficient of variation (CV%).

### Single molecule array (Simoa)

The development of quantitative methods and the measurement of the pathway proteins was performed on the automated Simoa HD-1 Analyzer platform (Quanterix©, Lexington, MA, USA). This instrument uses the same reagents as conventional ELISA but uses femtoliter-sized reaction chambers approximately 2 billion times smaller than conventional ELISA. This will result in a rapid buildup of fluorescence if a labeled protein is present which make it possible to detect single molecules. The instrument has previously been described in detail [13].

### Reagents

Capture antibodies tAKT and pAKT (DYC887B), tERK and pERK (DYC1230C), pp70s6k (DYC896) (R&D Systems, Minneapolis, MN, USA) and cyclin d (ab218793, Abcam, Cambridge, UK) were covalently attached by standard carbodiimide coupling chemistry to carboxylated paramagnetic beads (Quanterix). The biotinylated detector antibodies and the calibrators were tAKT (DYC1775), pAKT (DYC887B), tERK (DYC1230C), pERK (DYC1018B) (R&D Systems), pp70s6k (DYC896) and cyclin d (ab218793) (abcam). Streptavidin-β-galactosidase (SβG), enzyme substrate resorufin-β-D-galactopyranoside (RGP) and all consumables including wash buffers, cuvettes, disposable tips, and discs were from Quanterix.

### Simoa protocol

The six analyses were developed as single-plex assays using a Simoa 2-step assay for tAKT and pAKT and a 3-step assay for tERK, pERK, pp70s6k and cyclin d. Before running the following reagents are prepared; capture beads (1/3) are mixed with helper beads (2/3) in bead diluent buffer (Quanterix) and diluted to a final concentration of 2.0*10^7^ beads/ml. The biotinylated detector antibodies are diluted in sample/detector diluent (Quanterix) to final concentrations of 0.2 mg/L for tAKT and pAKT, 0.1 mg/L for tERK, pERK, pp70s6k and cyclin d.

The SβG is diluted in SβG diluent (Quanterix) to 150 pM. After loading the prepared reagents and consumables, the calibrators are prepared in diluent A (Quanterix). The samples and controls are diluted 30-fold in diluent A and loaded onto the instrument in a 96-well microtiter plate. The calibrators and the controls are run in duplicates and the samples are single determinations. The following steps are performed by the instrument. For the 2-step assay, 25 µl of capture bead is pipetted into a cuvette together with 100 µl of sample, control or calibrator and 50 µl of biotinylated detection antibody. An incubation step is performed for 30 minutes and the beads are then magnetically separated and washed. For the 3-step assay, 25 µl of capture bead is pipetted into a cuvette together with 100 µl of sample, control or calibrator and incubated for 40 minutes. The beads are then washed and 100 µl of detection antibody is added and an incubation step is performed for 5 minutes followed by washing the beads. The following steps are identical for both the 2-step and 3-step assays. 100 µl of SβG is added to the cuvette by the instrument and an incubation step is performed for 5 minutes. The beads are then separated magnetically and washed following the addition of RGP substrate. The bead substrate mixture is then loaded on to the Simoa disc containing an array of 216,000 micro-wells and sealed with oil. If protein has been captured and labeled, the SβG hydrolyze the RGP substrate into a fluorescent product that can be measured. At low concentrations of proteins, beads carry either zero or low numbers of enzymes and protein concentration is quantified by counting the presence of “on” or “off” bead (digital). At higher concentration of protein, each bead carries multiple enzymes and the total fluorescence signal is proportional to the amount of protein in the sample (analog). Both the digital and analog calculations use the unit “average number of enzyme per bead (AEB)”. The concentrations of protein in the unknown samples are interpolated from the calibrator curves obtained by 4 parameter logistic regression fitting.

### Mutation analysis

The mutational statuses of PIK3CA, BRAF and KRAS mutations were investigated in the cancer tissue using droplet digital polymerase chain reaction (ddPCR). The method has been described in detail in CEB Thomsen et al. [14]. The most frequent KRAS and BRAF mutations were investigated (KRAS G12D, G12V and G13D and BRAF V600E). If negative for these mutations, samples were analysed for 14 KRAS mutations in codon 12, 13, 61, 117 and 146 and 9 NRAS mutations in codons 12, 13 and 61. These 27 KRAS and NRAS mutations were selected based on the literature [7;15] and cover mutations found in more than 0.2% of colorectal cancers. All samples were analysed for the four most common PIK3CA mutations (E542K, E545K, H1047R and H1047L). Of the 176 patients used in this study, 58 patients were Wt for all investigated mutations (33%). Patients with BRAF mutations (n=53, 30%), KRAS mutations (n=56, 32%), NRAS mutations (n=4, 2%) and PIK3CA mutations (n=18, 10%). Patients with mutual mutations for PIK3CA and KRAS or BRAF (n=14, 8%). One patient had no mutational data.

### Statistical methods

Data were evaluated using NCSS software version 2007 (Kaysville, UT, USA) using the following statistical analyses: Wilcoxon Signed Ranks test, Mann-Whitney U-test, Spearman’s ρ, Kaplan-Meier log-rank test. For all analyses, a p-value < 0.05 was considered statistically significant.

## Results

### Setting up tERK, pERK, tAKT, pAKT, cyclin d and pp70s6k on the Simoa as single-plex assays

Figure 2 shows the calibrator curves for tERK, pERK, tAKT, pAKT, cyclin d and pp70s6k assays on the Simoa. The concentrations of the calibrators range from 0 to 2000 pg/ml for pAKT, 0 to 3000 pg/ml for cyclin d and 0 to 5000 pg/ml for tAKT, tERK, pERK and pp70s6k. Comparing the Simoa assays with traditional immunoassays an increase in sensitivity from 8-fold to 250-fold was achieved. For each assay, concentrations of the detection antibody and SβG have been optimized together with testing different sample/calibrator diluents to minimize matrix effects. The final parameters used are described in the materials and methods.

**Fig 2.**
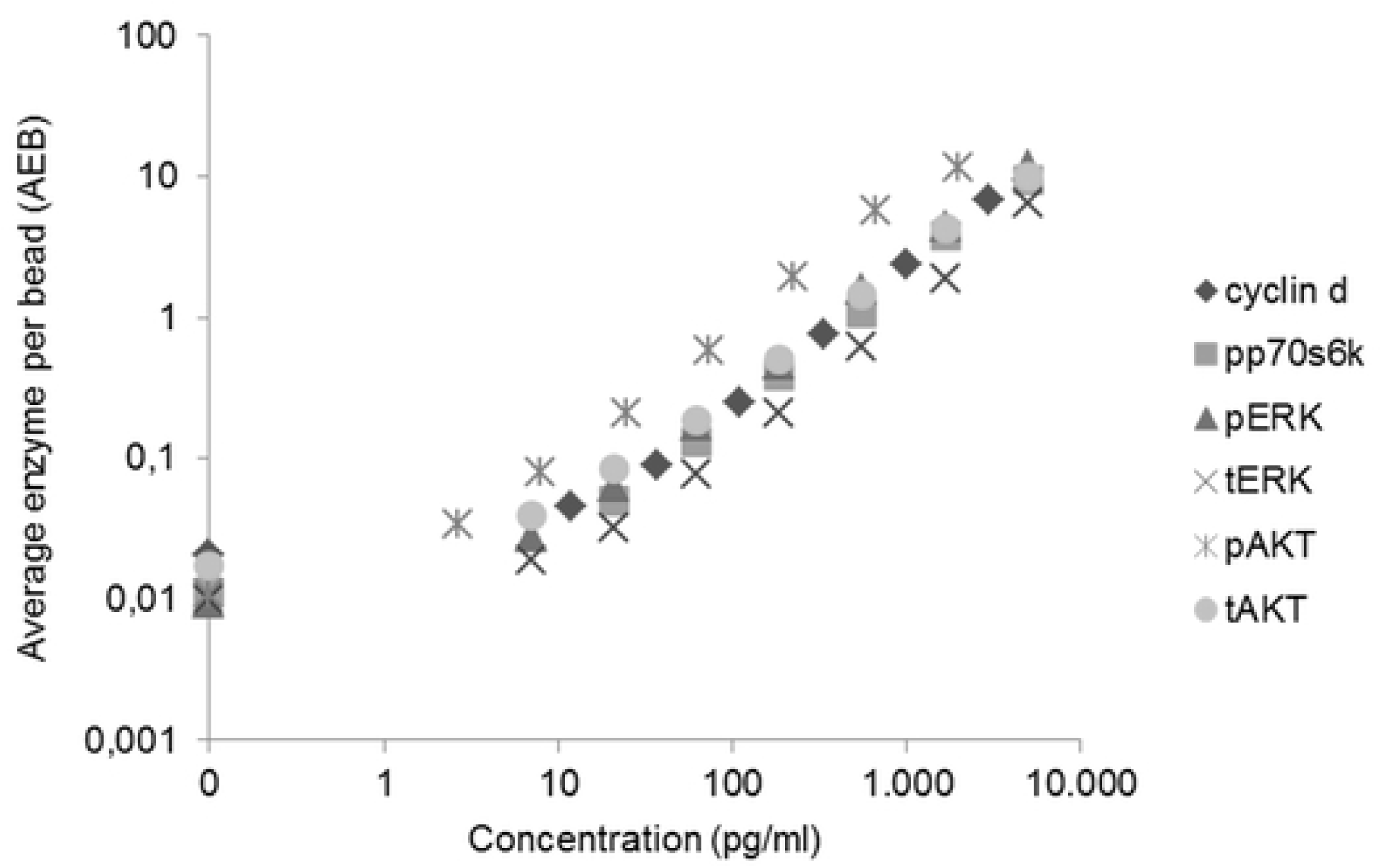
pERK, tERK, pAKT, tAKT, pp70s6k and cyclin d calibrator curves using Simoa. The average number of enzyme per bead (AEB) against concentration is shown. The calibrators (n=8) were run in duplicate and the mean value for each point of the calibrators is shown.

### Validating the single-plex assays

To study the matrix effects in the tissue lysate, samples were diluted in sample diluent ranging between 4 and 256 fold (4, 8, 16, 32, 64, 128, 256). The tissue samples show acceptable linearity between a dilution of 30 and 128 fold in all assays. In order to overcome matrix effects a 30-fold dilution of the samples was used. Limit of detection (LOD) was determined using 3 standard deviations (SD) from the background. Sample diluent was included at least 8 times over several days and the mean LOD was estimated (Table 1). For determining the intra-assay CV%, controls were analyzed in replicates of at least 6 in one assay and the total CV% was calculated from runs from different days (Table 1).

**Table 1.**
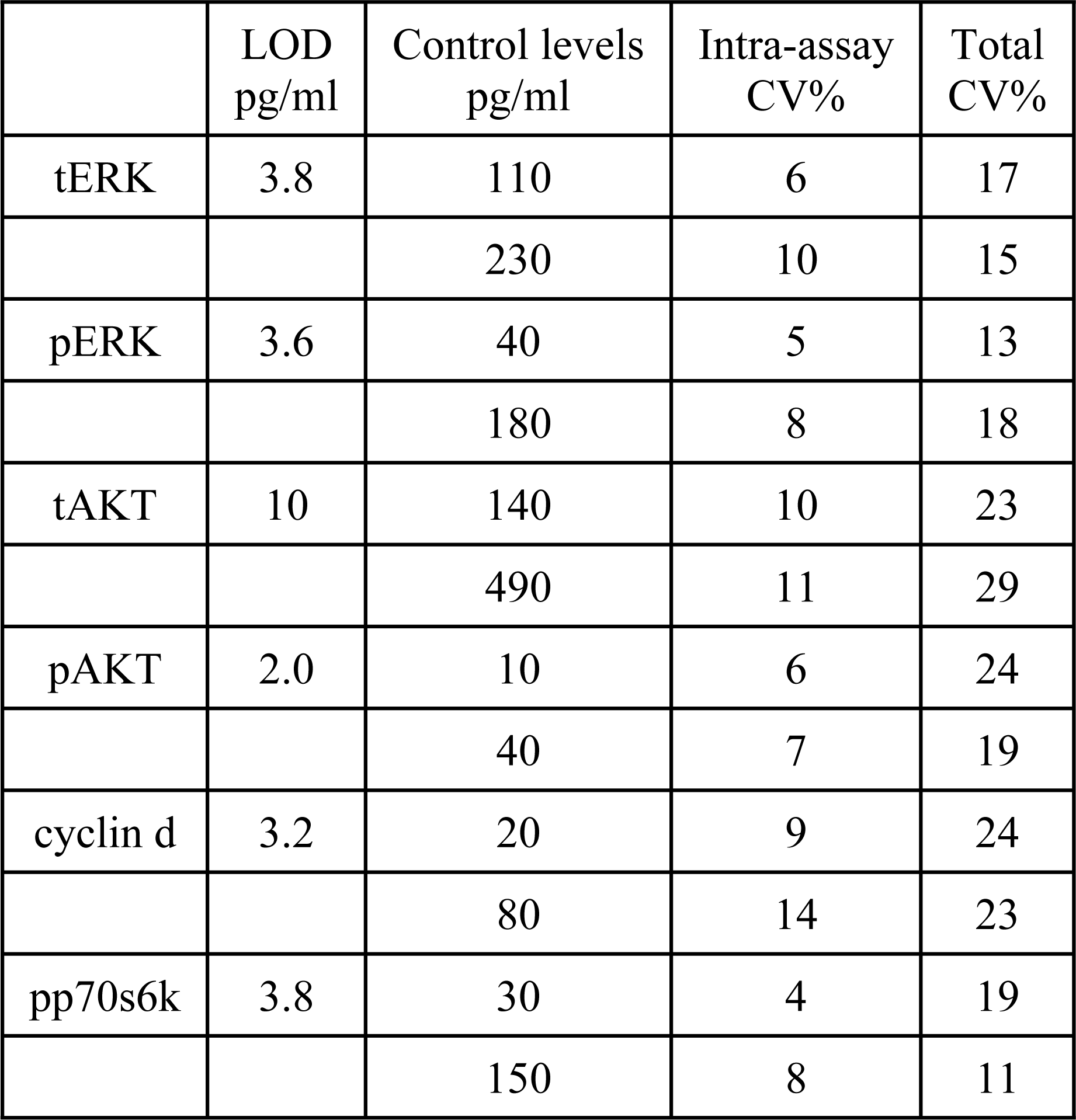
Performance of the assays.

### Autologous reference tissue and cancer tissue

tERK, pERK, tAKT, pAKT, cyclin d and pp70s6k were measured in both autologous reference tissue and colon cancer tissue (Fig 3 and Table 2). Both pERK and tERK were found to be down-regulated in cancer tissue (p= 0.001) while pAKT, tAKT, cyclin d and pp70s6k showed no differences between the tissues. Testing for variance differences between the tissue groups using Modified-Levene Equal Variance Test showed significant differences (p< 0.0001) for tAKT, pAKT, tERK and cyclin d.

**Fig 3.**
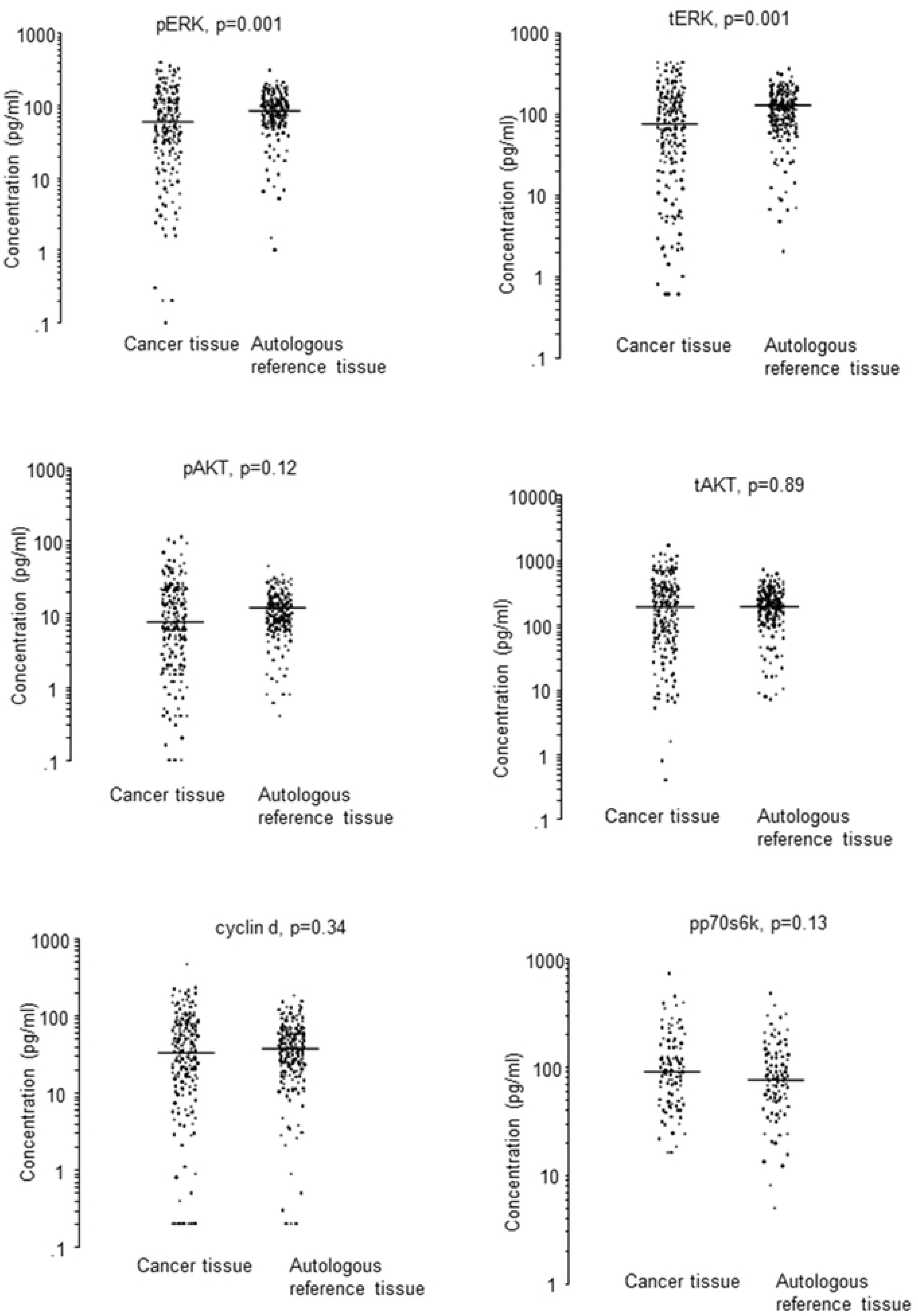
Pathway protein concentrations in cancer tissue and autologous reference tissue. The horizontal lines demonstrate the median values.

**Table 2.** Pathway protein concentrations in autologous reference and cancer tissue.

|  | Autologous reference tissue<br>Median (range) pg/ml | Cancer tissue<br>Median (range) pg/ml |
| --- | --- | --- |
| tERK | 111 (2-349) | 75 (0.6-421) |
| pERK | 83 (1-8174) | 61 (0.1-391) |
| tAKT | 202 (7-721) | 199 (0.4-1674) |
| pAKT | 11 (0.4-46) | 8 (0.1-115) |
| cyclin d | 37 (0.2-183) | 33 (0.2-470) |
| pp70s6k | 78 (5-480) | 93 (16-739) |

The median and range are shown for each pathway protein.

### Correlations between the pathway proteins

Associations between the pathway proteins in autologous reference tissue or cancer tissue were investigated. The Spearman’s rank correlation coefficients are shown in Table 3. In the autologous reference tissue the correlation between tAKT and pp70s6k was statistically significant (p= 0.0003). All other cases also showed statistically significant correlations (p<0.000001). Furthermore significant correlations were found between cancer tissue and autologous reference tissue regarding tAKT (p=0.003) and cyclin d (p=0.03).

**Table 3.** Correlations between the pathway proteins.

| R values | pERK | tAKT | pAKT | cyclin d | pp70s6k |
| --- | --- | --- | --- | --- | --- |
| Cancer tissue |  |  |  |  |  |
| tERK | 0.91 | 0.80 | 0.85 | 0.81 | 0.50 |
| pERK | - | 0.85 | 0.90 | 0.86 | 0.54 |
| tAKT | - | - | 0.93 | 0.81 | 0.54 |
| pAKT | - | - | - | 0.88 | 0.58 |
| cyclin d | - | - | - | - | 0.61 |
| Autologous reference tissue |  |  |  |  |  |
| tERK | 0.93 | 0.74 | 0.74 | 0.69 | 0.56 |
| pERK | - | 0.75 | 0.76 | 0.67 | 0.53 |
| tAKT | - | - | 0.78 | 0.58 | 0.37 |
| pAKT | - | - | - | 0.66 | 0.53 |
| cyclin d | - | - | - | - | 0.77 |

R values were calculated using Spearman's rank.

### Pathway proteins and mutational status

The concentrations of tERK, pERK, tAKT, pAKT, cyclin d and pp70s6k in colon cancer tissue were compared with the mutational statuses Wt, KRAS, BRAF and PIK3CA. The median and range values for all proteins are shown in Table 4. Patients with a BRAF mutation had significant decreased concentrations of tERK (p=0.0004), pERK (p=0.0002), tAKT (p=0.0002), pAKT (p=0.00004) and cyclin d (p=0.001) as compared with Wt (Fig 4). Patients with a PIK3CA mutation had significant decreased concentrations of tAKT as compared to Wt (p=0.042). There were no significant differences between patients with a KRAS mutation and Wt patients for either pathway protein.

**Table 4.**
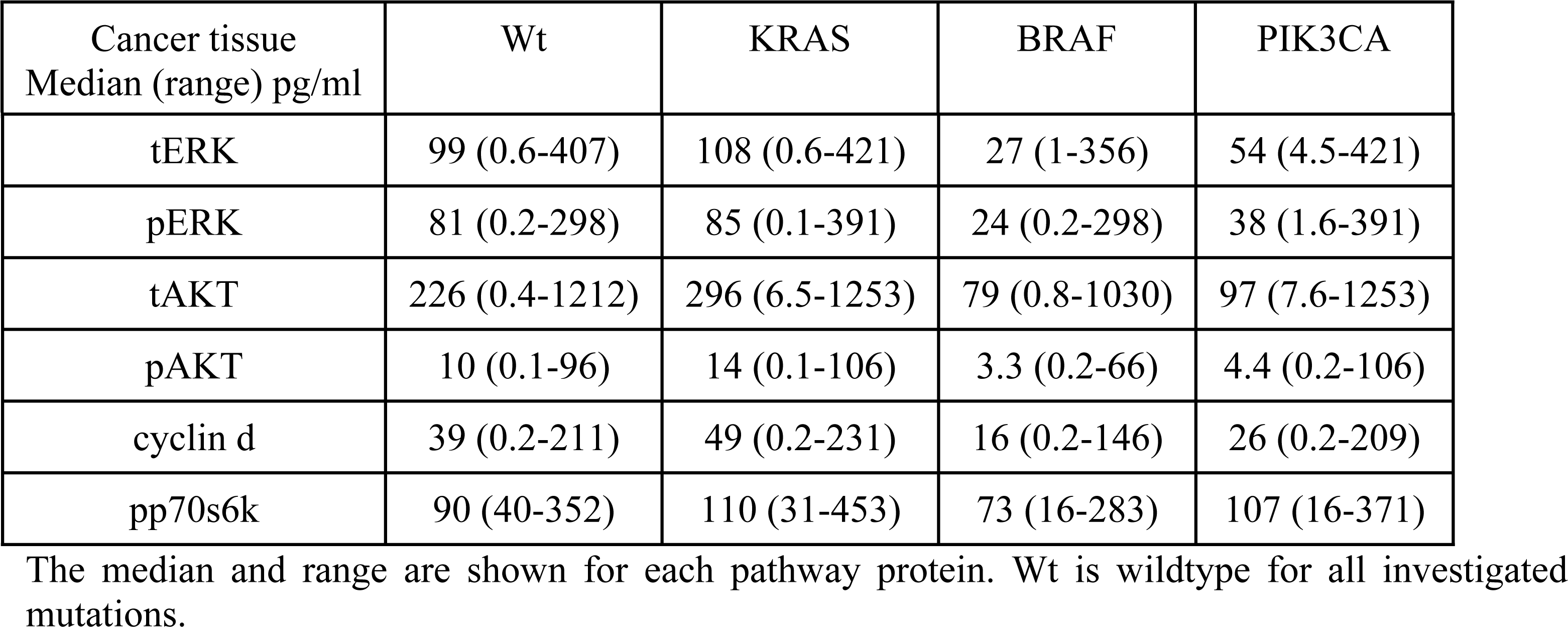
Pathway protein concentrations according to mutation status.

| Cancer tissue<br>Median (range) pg/ml | Wt | KRAS | BRAF | PIK3CA |
| --- | --- | --- | --- | --- |
| tERK | 99 (0.6-407) | 108 (0.6-421) | 27 (1-356) | 54 (4.5-421) |
| pERK | 81 (0.2-298) | 85 (0.1-391) | 24 (0.2-298) | 38 (1.6-391) |
| tAKT | 226 (0.4-1212) | 296 (6.5-1253) | 79 (0.8-1030) | 97 (7.6-1253) |
| pAKT | 10 (0.1-96) | 14 (0.1-106) | 3.3 (0.2-66) | 4.4 (0.2-106) |
| cyclin d | 39 (0.2-211) | 49 (0.2-231) | 16 (0.2-146) | 26 (0.2-209) |
| pp70s6k | 90 (40-352) | 110 (31-453) | 73 (16-283) | 107 (16-371) |

The median and range are shown for each pathway protein. Wt is wildtype for all investigated mutations.

**Fig 4.**
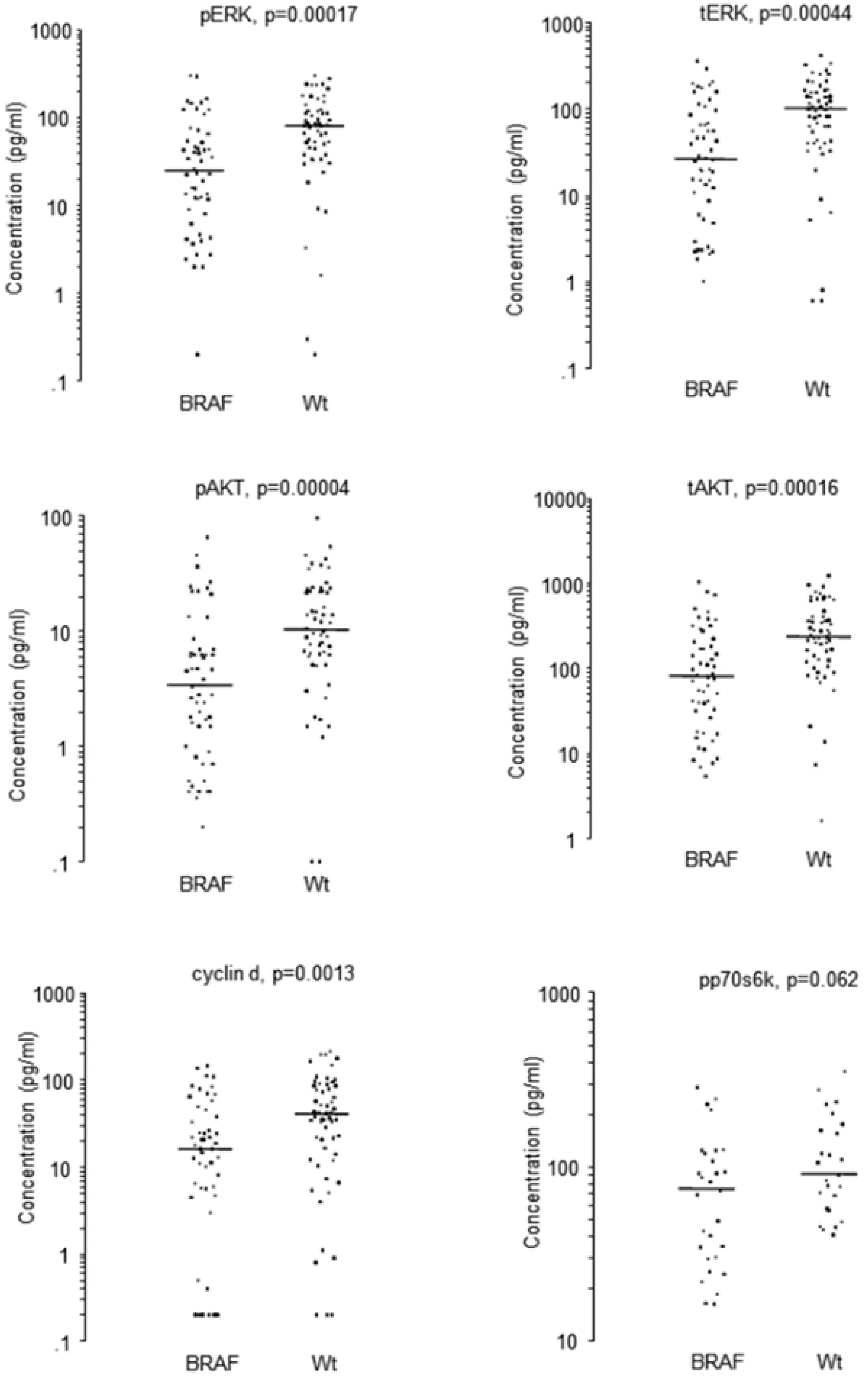
Pathway protein concentrations in cancer tissue with BRAF mutations or wt. The horizontal lines depict the median values.

### Clinical data

For each pathway protein the median concentration value was used as cut off and tested for its ability to distinguish between patients with different prognosis. Also the autologous reference tissue was used to establish cut off values for each pathway protein. There were no significant differences between patients with high concentrations and those with low concentrations using either discrimination cut off regarding disease free survival or overall survival.

Using the cut off values established from the autologous reference tissue and categorizing the cohort according to BRAF mutations a decreased overall survival was observed for patients with high levels of tERK (p=0.046), tAKT and pAKT (p=0.003) (Fig 5). Moreover patients with BRAF mutations showed decreased disease free survival for pERK (p=0.03), tAKT (p=0.019) and pAKT (p=0.034) (data not shown). Patients with KRAS mutations and low levels of pp70s6k demonstrated decreased overall survival (p=0.04) (Fig 5). Furthermore patients with KRAS mutations, stage 2 cancer and low concentrations of pp70s6k (p=0.003), cyclin d (p=0.045), tAKT (p=0.03), pAKT (0.03), tERK (p=0.04) or pERK (p=0.04) had inferior overall survival (Fig 6).

**Fig 5.**
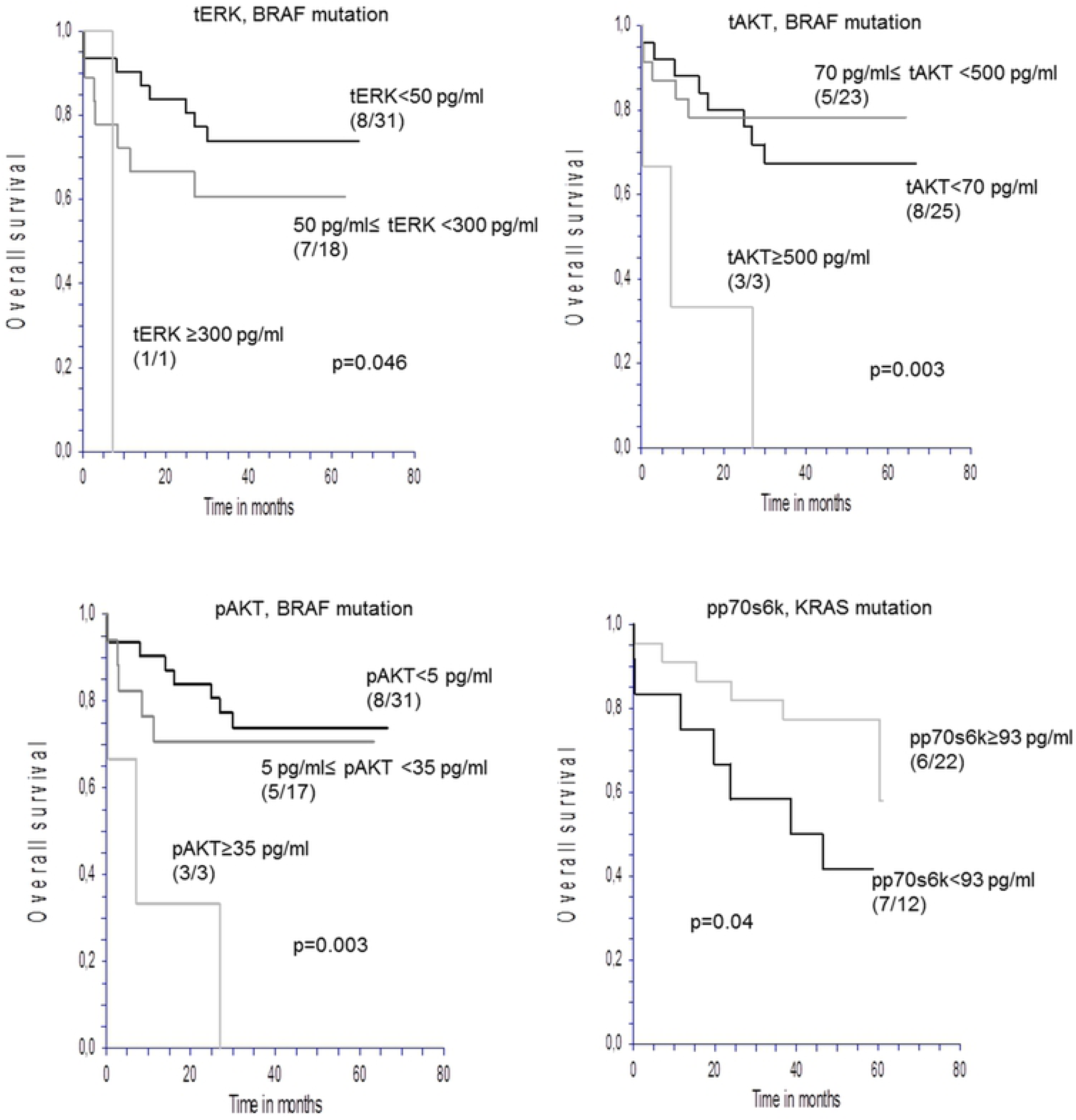
Overall survival in patients with BRAF or KRAS mutations. Kaplan-Meier curves. Numbers in parentheses indicate events/total number of patients.

**Fig 6.**
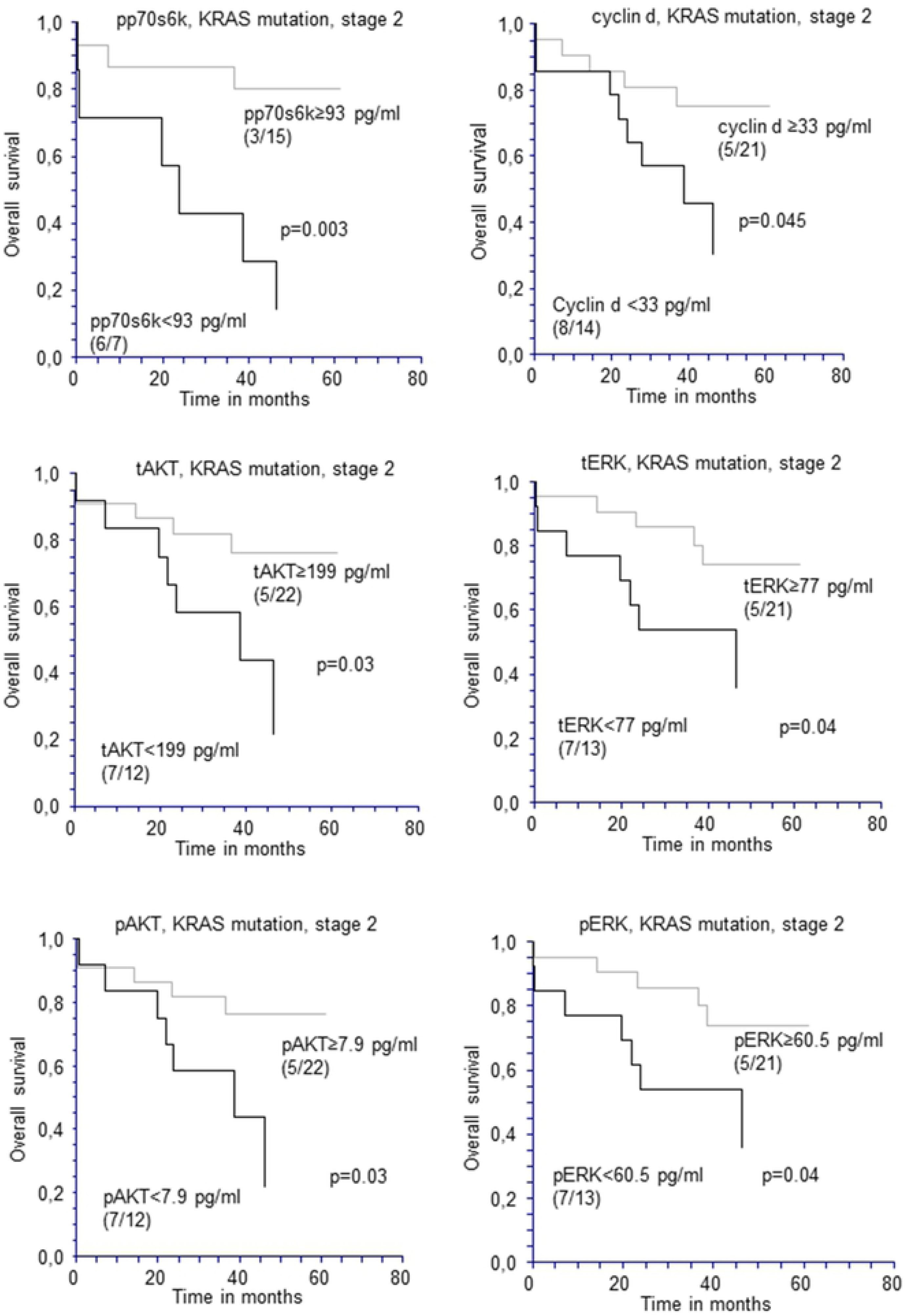
Overall survival in patients with stage 2 disease and KRAS mutations. Kaplan-Meier curves. Numbers in parentheses indicate events/total number of patients.

## Discussion

In this study quantitative methods were developed using Simoa technology for measuring AKT, ERK, cyclin d and pp70s6k in localized colon cancer tissue to investigate their correlation to prognosis and their relation to upstream mutations in the pathways.

The colon tissues were stored in RNAlater which might be a limitation of the study since the RNAlater solution contains a high level of ammonium sulfates which denature proteins. We therefore ensured that the developed assays were compatible with denatured proteins. Other studies have found that tissue preserved in RNAlater is suitable for ELISA-based methods [16;17].

PERK and tERK were found to be significantly down-regulated in cancer tissue as compared with autologous reference tissue which is also in agreement with the literature [18–22]. Furthermore, cancer tissue with a BRAF mutation demonstrated significant lower concentrations of pathway proteins as compared with Wt. We included 53 patients with a BRAF mutation while other studies using CRC patients have less than 12 patients with BRAF mutations included. The limited number of patients in these studies may be the reason for the divergent results they demonstrate [23–25].

No statistically significant correlations were found between ERK, AKT, pp70s6k or cyclin d and disease free survival or overall survival in the localized colon cancer patient cohort used in this study. However, stratifying according to mutation status, patients with BRAF mutations and high concentrations of ERK or AKT had low overall survival. Furthermore patients with KRAS mutations and low levels of pp70s6k had decreased overall survival and patients with KRAS mutations, stage 2 cancer and low concentrations of any of the measured pathway proteins demonstrated decreased overall survival. These results are based on a limited number of patients in each group and more patients are needed to document these findings. Studies on AKT or ERK activation have yielded variable results regarding survival. Malinowsky et al. showed that activation of AKT correlated with decreased survival, while Baba et al. showed that AKT activation was associated with a favorable outcome. Schmitz et al. found that the activation of ERK but not AKT predicted poor prognosis [25–27]. The majority of these studies included both colon- and rectal tumors and differences in prognostic value in the two groups may be possible. Also method differences, mutation data, sample sizes are some of the factors that might explain the divergent results.

Cancer tissues with genetic mutations result in a constitutive activation of the intracellular pathways. It is therefore plausible, that an increased activation and turn-over may lead to an over-production or consumption of native proteins as seen in the complement and coagulation pathway cascades. In this study there was an overall statistically significant correlation between pathway protein concentrations and mutational status. However, the change in pathway protein concentrations is too small to be used as screening indicators for mutations in practical medical use and our hypothesis could therefore not be confirmed. As in the complement and coagulation pathways the correct way to detect an increase in activity is to quantify not the native proteins but degradation or split products from the single intracellular pathway proteins. Therefore, we now aim to develop specific antibodies and methods to measure these degradation products, as previously done for complement C3d [28;29]. We believe this approach will increase both the prediction of mutations and survival.

## Acknowledgements

The authors thank the laboratory technologists Sara Egsgaard, Camilla Davidsen and Lone Karlsen Jensen, Department of Immunology and Biochemistry, for excellent technical assistance and continuous dedicated work.

